# Spatial modelling improves genomic evaluation in Tanzanian smallholder admixed dairy cattle

**DOI:** 10.1101/2025.03.01.640883

**Authors:** Isidore Houaga, Raphael Mrode, Julie Ojango, Chinyere Charlote Ekine-Dzivenu, Mwai Okeyo, Zabron Nziku, Athumani Nguluma, Eva Lavrenčič, Finn Lindgren, Ivan Pocrnic, Appolinaire Djikeng, Gregor Gorjanc

## Abstract

**Background:** Smallholder dairy production systems in low-and middle-income countries are characterised by large phenotypic variance due to diverse environmental effects, farming practices, and crossbreeding. Furthermore, small herds, low genetic connectedness, and limited data recording challenge accurate separation of environmental and genetic effect in such settings, limiting genetic improvement. Here, we evaluated the impact of modelling spatial variation between herds to address these challenges and improve the accuracy of genomic evaluation for Tanzanian smallholder dairy cattle.

**Results:** We analysed 19,375 test-day milk yield records of 1894 dairy cows from 1386 herds across four distinct geographical regions in Tanzania. The cows had 664,822 SNP marker genotypes after quality control and were highly admixed. We fitted a series of GBLUP models to evaluate the impact of modelling the herd effect and the spatial effect on. The herd effect was fitted as an independent random effect, while the spatial effect was fitted as a random effect with Euclidean distance-based Matérn covariance function. The models were compared based on: model fit; estimates of variance components and breeding values; correlations between the estimated contribution of breeding values, herd effect, and spatial effect to phenotype values; and the accuracy of phenotype prediction in cross-validation and forward validation. The results showed large differences in milk yield between and within regions, as well as significant variation due to the spatial effect, which were not fully captured by modelling the herd effect. The results also strongly indicate that a model with just the herd effect underestimated breeding values of animals in less favourable environments and overestimated breeding values of animals in more favourable environments.

**Conclusions:** This study demonstrated the challenge of achieving accurate genomic evaluation in smallholder settings. By leveraging spatial modelling we maximised the use of available data and improved the separation of genetic and environmental effects. Further work is required to improve smallholder genetic evaluations by understanding environmental and genetic processes that drive the large phenotypic variance in African smallholder setting.

## Background

This paper demonstrates how spatial modelling improves genomic prediction in a population of dairy cattle with large phenotypic variance due to diverse environmental effects, farming practices, and crossbreeding of smallholders in low-and middle-income countries (LMIC). The livestock sector plays an important role in improving the livelihoods of people in LMIC through income generation, food supply, improved nutrition, and consequently, health. However, its impact on the economy and contribution to the gross domestic product is generally below its potential and is highly varying between countries. In developed countries, the significant increase of milk yield in the dairy cattle sector has been due to selective breeding and herd management strategies [1]. For example, introduction of genomic selection increased genetic gains for milk, protein and fat yields for registered cows in the USA, respectively, from 50, 1.6, and 2.2 kg per year to 109, 4.1, and 6.0 kg per year [2]. In many LMICs, genetic improvement is still in its infancy, mainly due to the prolonged absence of a systematic approach to routinely collect performance data [3]. In the context of smallholder dairy production systems in LMIC and the associated lack of pedigree information, the use of genomic selection presents an even greater opportunity compared to advanced economies [4, 5].

In this sense, many efforts have been made to improve low productivity levels in LMIC smallholder dairy cattle, including the Africa Asia Dairy Genetic Gains project (AADGG; https://www.ilri.org/research/projects/aadgg). The AADGG has generated performance and genomic data on smallholder dairy cattle systems across East Africa, including Tanzania. For example, see [3, 4], and [6] reports on genetic evaluation of daily milk yield based on the AADGG data. These studies based their genetic evaluation on genomic best linear unbiased prediction (GBLUP) [7] and single-step GBLUP (ssGBLUP) [8].

Genetic evaluation separates genetic and environmental effects on an individual’s phenotype [9, 10]. This separation is important for related or unrelated individuals in shared or different environments. Practically, this separation is achieved by defining contemporary groups of individuals, which share an environment and estimating the effect of these groups on phenotypes. To achieve high accuracy, genetic evaluation requires substantial collection of phenotypic data, large, yet homogeneous, contemporary groups, and genetic connectedness between the contemporary groups. However, the African dairy production systems have small herds, low genetic connectedness between herds, and even low genetic relationships between animals within herds [3, 5]. It is challenging to achieve high accuracy of genetic evaluation in such conditions [11, 12].

The accuracy of genetic evaluation in breeding programmes with small herds is an important topic that has been studied with simulation and real data (see [12] for a recent summary of literature). Powel et al. [5] simulated a breeding programme similar to those in East Africa and studied the accuracy of standard genetic evaluation models in separating genetic effects from herd effects when herd sizes are very small. They found that modelling small herds (≤8 cows) as a random effect achieved higher accuracy than modelling them as a fixed effect. At larger herd sizes, the accuracy of fixed and random effect modelling converged. Costilla et al. [13] used the BayesR model [14] with herd effects as random for genetic evaluation of Indian smallholder dairy cattle. Similarly to [5], they found that the accuracy of genetic evaluation increased when herd size increased - up to about 5 cows and became constant thereafter.

Accounting for shared environmental conditions can enhance the separation of environmental effects from genetic effects. Nearby herds are likely to experience similar climate, soil, management, and even societal aspects related to farming, such as road infrastructure that facilitates trade, availability of agricultural schools, etc. In the absence of information on these environmental conditions and their complex interactions, we can attempt to account for the shared environment among herds by accounting for distance between them. Sæbø and Frigessi [15] analysed mastitis resistance across Norway with the Besag’s regional model [16, 17]. They estimated mean differences between farms within the regions of Norway without explicitly fitting herd effects. Chawala et al. [18] in their simulation study defined villages as a contemporary group to share information between farms within a smaller region. This was done in the context of small farms in Sub-Saharan Africa to enable accurate estimation without estimating the effects of small herds. Their approach however ignores environmental variation within villages, like Sæbø and Frigessi [15] approach ignores variation within regions. Mrode et al. [4] similarly modelled administrative region (ward) as a fixed effect, but also herd as a fixed effect despite the small herd size in the AADGG data. While these approaches model shared environmental conditions in some way, neither approach is sufficiently general. In human genetics, Heckerman et al. [19] explicitly addressed this issue by leveraging coordinates of the sampled individuals and used a coordinate-based model from the field of geostatistics [17, 20]. In these models, location is modelled as a random effect with distances between locations informing covariance between location effects. Selle et al. [11] and Cuyabano et al. [12] are examples of such approaches to model the herd effect in animal breeding. Selle et al. [11] used the computationally scalable approach from [21] with the general Matérn covariance function between herd locations and have shown that this approach significantly improved the accuracy of estimated breeding values (EBV) compared to random independent herd effects in simulated and real data. Cuyabano et al. [12] used exponential covariance function between herd locations of Hanwoo beef cattle and also observed increased accuracy compared to random independent herd effects. The exponential covariance function is a special case of the Matérn covariance function.

While Selle et al. [11] and Cuyabano et al. [12] clearly show that spatial modelling increases the accuracy of genetic evaluation, both studies were based on data from developed countries with good genetic connectedness. Selle et al. [11] tried to mimic small herds situation observed in LMIC by sampling animals from herds, but could not mimic the low genetic connectedness common across herds in LMIC due to the low use of artificial insemination. Cuyabano et al. [12] tested spatial model on data from 124 commercial beef cattle farms with the average herd size of 17 animals and more than 66% of farms having more than 5 animals, which is far from the LMICs smallholder context with 95% of farms with less than 3 dairy cows [3]. Testing spatial modelling for genetic evaluations is required for a challenging case such as LMIC data. This data has small herds, low genetic connectedness, and substantial components of genetic and environmental variation.

The aim of this study was therefore to evaluate the potential of spatial modelling for genetic evaluation of data from the AADGG programme in Tanzania with its georeferenced herds. We evaluated the impact of spatial modelling on estimates of variance components and breeding values, correlation analysis between models and their model components, and the accuracy of predicting milk yield phenotype. The estimated spatial effects showed strong regional differences, important impact on the EBV, and improved the accuracy of phenotype prediction.

## Methods

### Animals and genotype data

We analysed data from 1894 admixed cows involving East African indigenous breeds, mostly represented by N’dama and Small Eastern Africa Zebu, and 5 exotic *Bos taurus* dairy breeds (Ayrshire, British Friesian, Guernsey, Holstein, and Jersey) [4]. For this study, genotype data were available for 1894 cows registered in the AADGG programme in Tanzania from November 2016 to May 2020. The cows were genotyped with the GeneSeek Genomic Profiler Bovine 50K array with 47,843 SNP markers. The quality controls applied on the genotype data were: only SNP with a call rate of above 95% were kept, SNP markers with a minor allele frequency less than 0.01 were excluded, and only animals with genotypes for more than 90% of SNP markers were retained. After the quality control, the remaining 40,581 SNP markers were imputed to 664,822 SNP markers from the Illumina high density array using a reference population consisting of crossbred animals (*Bos indicus* × *Bos taurus*) from a previous project and several European purebreds, including Ayrshire, British Friesian, Guernsey, Holstein, and Jersey [22]. Given that the use of different breeds is not systematic, we refer to such crossbred animals as admixed. The breed composition of each cow in these data was previously estimated as the percentage of genome from exotic and indigenous populations using an admixture analysis [22]. The percentage of exotic genome for each cow was calculated as the sum of the estimated percentage contributions of each of the 5 exotic dairy breeds. Following Mrode et al. [4] four levels for the percentage of exotic genome were used: [100, 87.5], (87.5, 60], (60, 36], and (36, 0]. These classes were used to account for the effect of admixture in all subsequent analyses.

The cows were from 1386 herds. All the herds were geographically referenced using Global Positioning System (GPS). Herd coordinates were received as latitude and longitude degrees based on the EPSG:4326 geographic coordinate system and the World Geodetic global reference system 1984 (datum WGS84) [23] We projected the coordinates for Tanzania and expressed them in kilometres using the EPSG:21035 geographic coordinate system. The herds were located within 72 wards, 9 districts, and 6 regions of Tanzania. A ward is an administrative unit comprising several villages and was defined after mapping the GPS coordinates of the herds to the Tanzania administrative level 3 shapefile (https://data.humdata.org/dataset/cod-ab-tza?). As shown in Figure 1, the herds were clustered in 4 regions defined as follows: North-Central (latitude *>* 9500), North-East (longitude *>* 1500 & latitude *<* 9500), South-Central (longitude *>* 1300 & latitude *<* 9200), and South-West (longitude *<* 1300 & latitude *<* 9200). The figure also shows a triangulation mesh that is used as a computational approach to spatial modelling with Matérn covariance function, described in the statistical modelling sub-section of this manuscript. The mesh is denser in the area of observed locations and less dense beyond the regions; and the meshed area spans beyond the borders of Tanzania to avoid edge effects.

**Figure 1.**
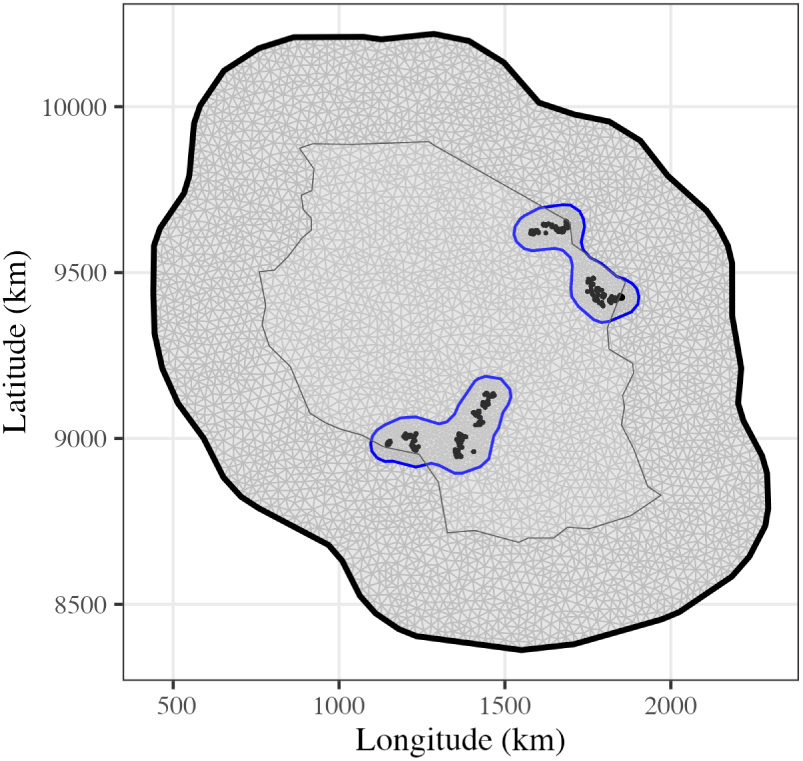
Sampling sites of the AADGG project in Tanzania from November 2016 to May 2020 with 1386 herds clustered in North-Central, North-East, South- Central, and South-West region (the gray triangulated mesh is described in the text)

We conducted the principal components analysis to understand the genetic structure of the studied population. The PCA was performed using the prcomp function of the R statistical software [24] We further conducted a multivariate analysis of variance (MANOVA) to test for difference in SNP genotype variation between the defined exotic genome percentage levels as well as regions.

### Performance data

Performance data consisted of test-day milk yield extracted from the AADGG database at the International Livestock Research Institute (https://www.adgg.ilri.org/uat/auth/auth/login). The initial data consisted of 19,538 test-day milk yield records. The data were subjected to several filters: daily milk yield values in the range between 1 and 45 litres; age at first calving of at least 18 months; and days in milk (DIM) in the range between 4 and 500 days. The DIM upper limit was based on a preliminary analysis that showed a reduction in heritability for milk yield with DIM greater than 500 [4]. Parity was represented in the initial data with levels from 1 to 9. However, we fitted parity effect with only 3 levels: parity 1 and parity 2 separately, and parities 3+ pooled into the third level. Additional metadata included calving season with 2 levels per year (dry or wet), where January to June is the dry season. After data editing, we were left with 19,375 test-day records for 1894 cows from 1386 herds for further analyses. The herd size ranged from 1 to 9 with more than 71% of herds having a single cow and more than 93% herds with less than 3 cows. For downstream statistical modelling, milk yield and age at calving were standardised to mean zero and unit variance. Hence, all reported estimates related to milk production are expressed in units of phenotypic standard deviation or variance.

### Statistical modelling

Following Mrode et al. [4], the following baseline (G) model was fitted to the observed phenotype *y_i_* of individual *i*:

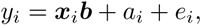

where ***b*** is a vector of fixed effects, consisting of the intercept, 4 levels for the percentage of exotic genome, calving year-season, test year-month, age effect nested within parity level, and lactation curves modelled by Legendre polynomials of order 2 nested within parity level, *a_i_* is the additive genetic effect (breeding values), and *e_i_* is the residual effect with *e_i_* ∼ *N* (0, ***I****σ_g_*^2^) where *σ_e_*^2^ is residual variance. We modelled the breeding values as ***a*** ∼ *N* (**0**, ***G****σ_g_*^2^), where ***G*** is the VanRaden type 1 genomic relationship matrix [7] and *σ*^2^ is genomic variance.

We then extended the G model in three ways. The GH model included the herd effect assuming independent and identically distributed herd effects, *h_i_* ∼ *N* (0, ***I****σ_h_*^2^), where *σ*^2^ is herd variance. The GS model included the herd location effects assuming Matérn covariance between locations, ***s*** ∼ *N* (**0***, M* (***S****, ρ, σ_s_*^2^)), where ***S*** is the Euclidean distance matrix between locations and the two hyper-parameters model the variation of location effects (*σ_s_*^2^) and the range of spatial correlation (*ρ*); defined as the distance where correlation between two points is near 0.1 (see [11, 21] for details). The GHS model combined all these effects into single model.

Preliminary analyses indicated challenges in fitting the permanent environmental effect. We associated this challenge to the data structure with a very high proportion of herds with a single cow (71%). With such data structure, it is difficult to correctly separate breeding value, permanent environmental, and herd effects. For comparison, we fitted an alternative set of models assuming independent and identically distributed permanent environmental effects, *p_i_* ∼ *N* (0, ***I****σ_p_*^2^). These corresponding models with permanent environmental effects are GP, GPH, GPS, and GPHS.

We used the Bayesian numerical approximation procedure known as the Integrated Nested Laplace Approximation (INLA) introduced by Rue et al. [25] with further developments and implementations in the *R-INLA* and *inlabru* R packages [26, 27, 28]. These two packages enable the use of a computationally scalable approach for Bayesian spatial modelling in the context of a linear mixed model [21]. A full Bayesian analysis requires prior distributions for all model parameters. We assumed ***β*** ∼ *N* (0*, σ_β_*^2^) with *σ_β_*^2^ = 1000 for the intercept and fixed effects. Following the previous study by Selle et al. [11], we assumed penalised complexity priors for the spatial effect hyper-parameters [29], and default R-INLA priors for other variance parameters. Lastly, because the studied herds were from only a few parts of Tanzania, we report in the results an estimate of *σ*^2^ as a model parameter and as the “realised” spatial variance at our herd locations by sampling 1000 realisations from the posterior distribution of spatial effects projected to herd locations by using the ‘inlabru::generate()‘ function, calculating the variance of the projected spatial effects for each sample, and summarising the resulting posterior distribution of the effects.

### Model evaluation

We evaluated the models with model fit statistics, correlation analysis between models and their model components, and cross-validation and forward validation of phenotype prediction. We used the deviance information criterion (DIC) [30] to compare the fit of the models, with lower values indicating a better model fit to the data. The correlation analysis first merged into one table EBVs from different models as well as estimated herd and spatial effects corresponding to animal’s herds. Then we correlated these merged estimates, which we refer to as the correlation between the estimated contribution of breeding value, herd, and spatial effects to animal’s phenotypes. To this end, we have constructed and correlated estimates of phenotypes (via a linear predictor) based on each of the estimated breeding values, herds, and spatial effects. Finally, accuracy of phenotype prediction between models was assessed with cross-validation and forward validation. The cross-validation involved the exclusion of all phenotypes for each level of cows of certain percentage of exotic genome or region and then predicting all the excluded phenotypes. Thus, 7466 records (731 cows), 8149 records (770 cows), 3058 records (309 cows), and 702 records (84 cows) were excluded respectively for the [100, 87.5], (87.5, 60], (60, 36], and (36, 0] percentage of exotic genome For regional cross-validation, 4547 records (437 cows), 8482 records (775 cows), 3674 (414 cows), and 2672 (268 cows) were excluded respectively for the North-East, North-Central, South-Central and, South-West region. The forward validation was implemented by excluding 1424 records from 146 cows born in 2016 and 2017.

## Results

In the results, we first present the summary statistics for the milk yield data, followed by the multivariate analysis of genomic data with the goal of understanding the genetic and spatial structure of the data. The main part of the results show the estimates of variance components with different models, including models with spatial effects and their correlations with the EBV. We close the results section with an accuracy assessment of the tested models to predict observed phenotypes.

### Summary statistics of milk yield

The summary statistic of milk yield across different levels of percentage of exotic genome and regions is presented in Table 1. The overall mean (±SD) of test-day milk yield was 8.3±4.3 L. The mean increased with parity as expected. The mean (±SD) milk yield for cows with [100, 87.5], (87.5, 60], (60, 36], and (36, 0] percentage of exotic genome was respectively 9.6±4.2, 8.2±4.2, 6.2±3.5, and 4.5±2.6 L. This indicates an increase in milk yield with an increase in the percentage of exotic genomes. The South-West region had the highest mean milk yield (12.4±4.4 L), while the North-East region had the lowest mean milk yield (6.7±3.9 L). The North-Central and South-Central regions had intermediate mean milk yield of ∼8 L.

**Table 1:** Mean and standard deviation (SD) of the test-day milk yield (L) across key factor levels.

| Variable | Number of cows | Number of test day records | Mean ± SD |
| --- | --- | --- | --- |
| Overall | 1894 | 19375 | $8.3 \pm 4.3$ |
| Parity |  |  |  |
| 1 | 1024 | 5883 | $7.3 \pm 4.1$ |
| 2 | 1206 | 7018 | $8.2 \pm 4.1$ |
| 3+ | 777 | 6474 | $9.2 \pm 4.4$ |
| Percentage of exotic genome |  |  |  |
| [100, 87.5] | 731 | 7466 | $9.6 \pm 4.2$ |
| (87.5, 60] | 770 | 8149 | $8.2 \pm 4.2$ |
| (60, 36] | 309 | 3058 | $6.2 \pm 3.5$ |
| (36, 0] | 84 | 702 | $4.5 \pm 2.6$ |
| Region |  |  |  |
| North-East | 437 | 4547 | $6.7 \pm 3.9$ |
| North-Central | 775 | 8482 | $8.0 \pm 3.9$ |
| South-Central | 414 | 3674 | $8.0 \pm 3.6$ |
| South-West | 268 | 2672 | $12.4 \pm 4.4$ |

The variation of daily milk yield across the levels of percentage of exotic genome and regions is illustrated in Figure 2. The test-day milk yield varies substantially, and as seen in Table 1, milk yield decreases with decreasing percentage of exotic genome. However, the North-Central and South-West regions had more animals with higher percentage of exotic genome ((87.5, 60] and [100, 87.5]), while the North-East and South-Central regions had comparatively more animals with lower percentage of exotic genome (36, 0] and (60, 36]).

**Figure 2.**
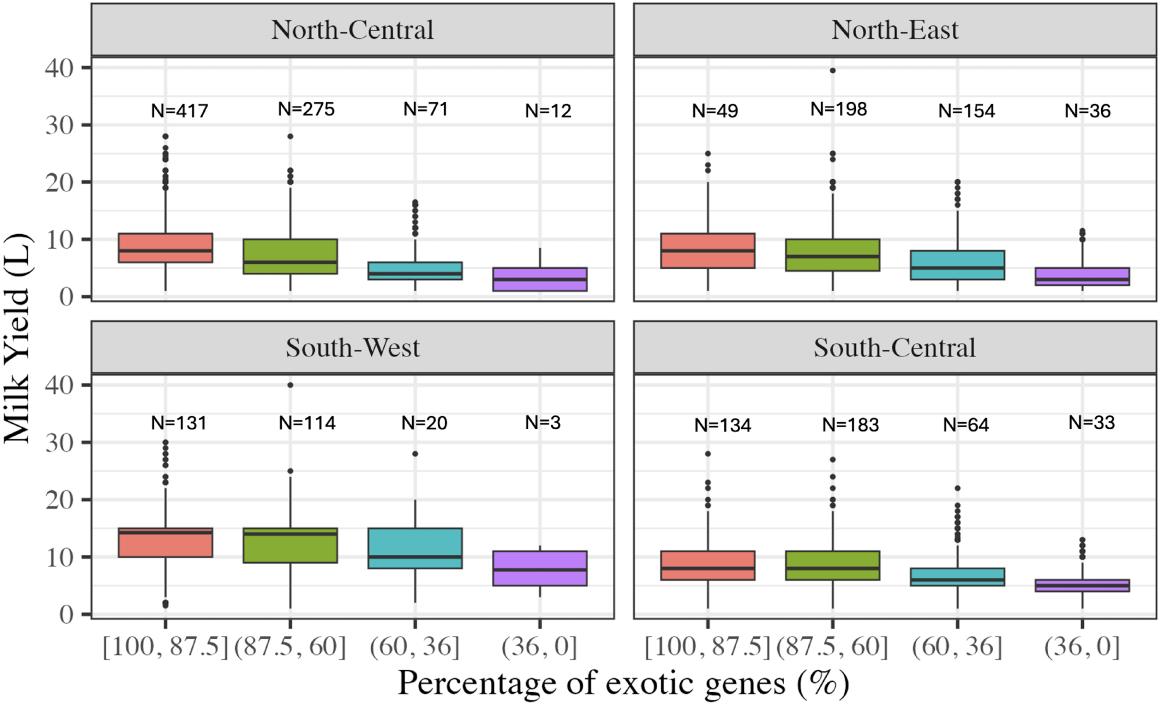
Boxplot of milk yield and number (N) of cows by the level of percentage of exotic genome and region

### Genomic and spatial population structure

To understand the genetic and spatial structure of the data, we performed two multivariate analyses. First, we estimated principal components to analyse the genetic structure of the studied population. The projection of SNP genotypes onto the first 3 principal components showed a substantial overlap between the levels of percentage of exotic genome and regions (Figure 3). To test for the difference in SNP genotype variation between these levels, we used Multivariate Analysis of Variance (MANOVA, Table 2). The first principal component differed significantly with the levels of percentage of exotic genome (*p <* 0.001), which was expected due to the high admixture in the population studied, but this only explained 3.1% variation in SNP genotypes. The decreasing value for the first principal component aligned with the decreasing level of percentage of exotic genome. The second and third principal components did not differ significantly with the levels of percentage of exotic genome, but did differ between regions (Table 2). Although not significant, North-East region had a lower mean for the first principal component compared to other regions, indicating less exotic admixture compared to other regions. The second and third principal components differed significantly between regions, indicating a different distribution of alleles across regions, although these two components collectively explained less than 1.3% variation in SNP genotypes.

**Figure 3.**
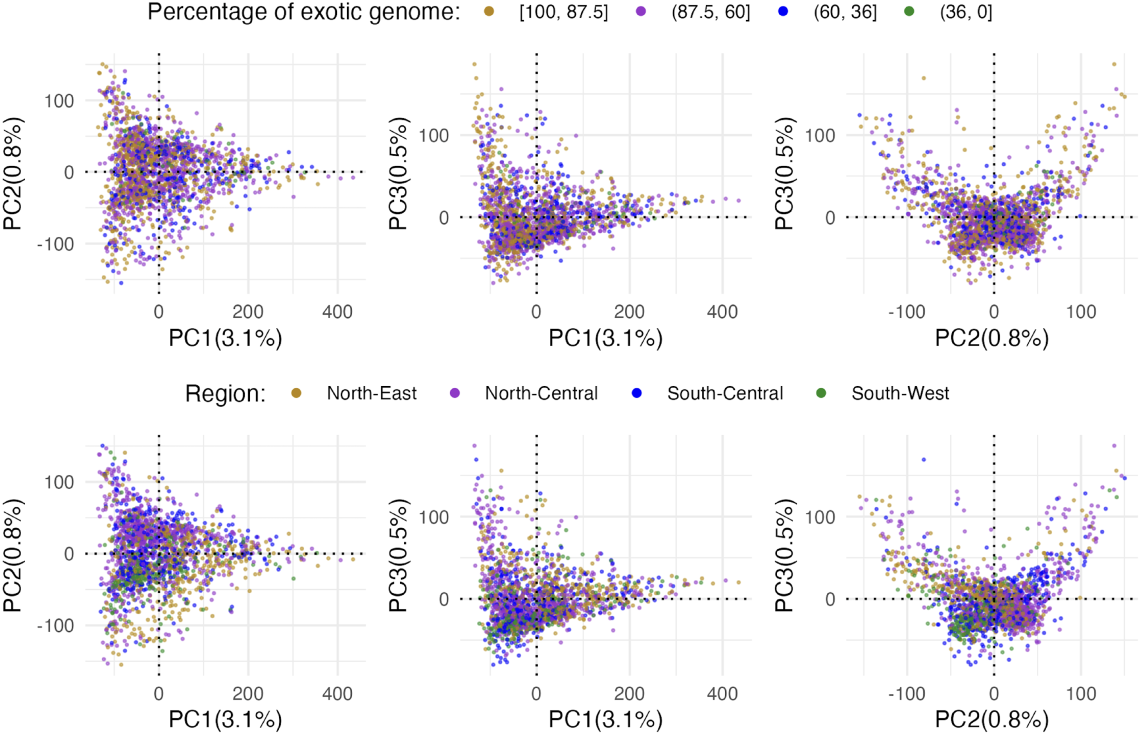
Projection of SNP genotypes onto the first 3 principal components coloured by the level of percentage of exotic genome (top) and region (bottom)

**Table 2:** Multivariate analysis of variance for principal components by the level of percentage of exotic genome and region - values shown are the mean ± Standard Error of principal components.

| Principal component<br>(variance explained) | Percentage of exotic genome |  |  |  | p-value |
| --- | --- | --- | --- | --- | --- |
|  | [100, 87.5]<br>(n=731) | (87.5, 60]<br>(n=770) | (60, 36]<br>(n=309) | (36, 0]<br>(n=84) |  |
| PC1 (3.1%) | 10.9±3.2 | -15.2±4.5 | -19.9±5.9 | -27.9±10.0 | 0.0001 |
| PC2 (0.8%) | -0.8±1.7 | 2.0±2.4 | 0.8±3.1 | -3.8±5.3 | 0.6658 |
| PC3 (0.5%) | 0.8±1.3 | -1.9±1.7 | -0.0±2.3 | -1.2±3.9 | 0.7028 |
| Principal component<br>(variance explained) | Region |  |  |  | p-value |
|  | North-East<br>(n=437) | North-Central<br>(n=775) | South-Central<br>(n=414) | South-West<br>(n=268) |  |
| PC1 (3.1%) | -7.8±4.1 | 12.4±5.2 | 7.9±5.9 | 9.1±6.7 | 0.1287 |
| PC2 (0.8%) | 1.3±2.1 | -6.3±2.7 | -2.3±3.1 | 11.9±3.5 | 0.0000 |
| PC3 (0.5%) | -0.9±1.6 | 2.6±2.0 | 2.7±2.3 | -5.7±2.6 | 0.0038 |

### Spatial effect

Results from statistical models that included the spatial effects (GS and GHS) have successfully captured the spatially dependent structure of the data. The estimated spatial effect from the GHS model is shown in Figure 4. The left panel shows the posterior mean and the right panel shows the posterior standard deviation (uncertainty) of the estimated spatial effect across the studied country. Non-zero estimates of the spatial effect aligned with the four regions of herds in the data; two with predominantly negative mean effect in the North-East and South-Central regions of Tanzania, and two with predominantly positive mean effect in the North-Central and South-West regions of Tanzania. The magnitude of the estimated mean spatial effects ranged from −2.5 to 1.5 (a range of 4) phenotypic standard deviation across Tanzania. These estimates align with the mean milk yield in Table 1. Since the herds clustered in the four regions, the rest of the country had an estimated spatial effect close to zero (shrinkage). Relatedly, the uncertainty (as measured by the posterior standard deviation) of the estimated spatial field was lower in the four regions where phenotypic data was collected than elsewhere. The estimated spatial effect from the GS model was similar, but not identical, to the estimates from the GHS model (Supplementary 1).

**Figure 4.**
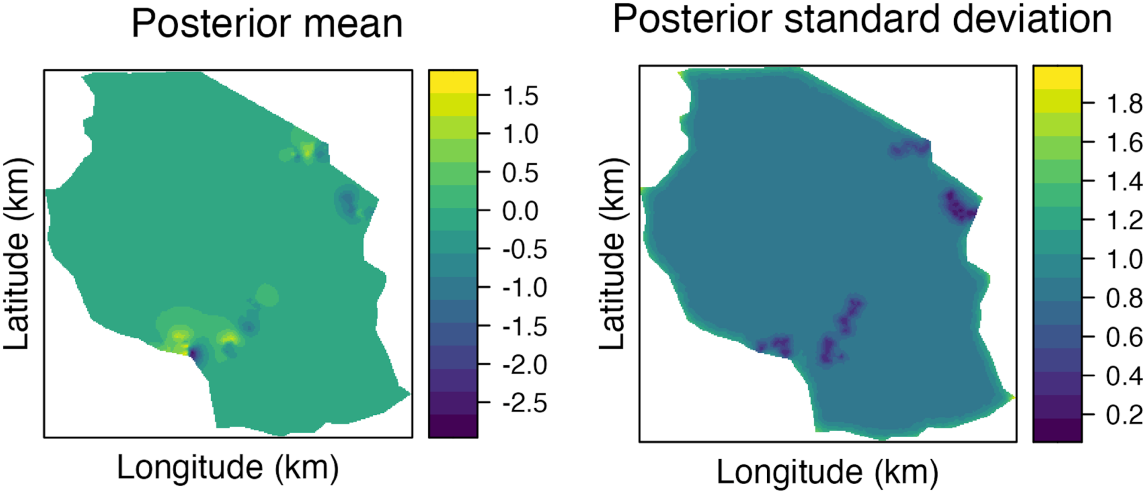
Posterior mean (left) and standard deviation (right) of the spatial effects (in units of phenotypic standard deviation) from the GHS model

### Model fit and variance components

We evaluated model fit to the data with DIC (Table 3). The GHS (DIC=32596) and GH (DIC=32650) models had the best fits given the model complexity, followed by G (DIC=32716) and GS (DIC=32735) models. DIC differences between these models indicate that modelling both spatial and herd effects jointly is important, and that modelling just spatial effect is even worse than not modelling either of these effects.

**Table 3:** Estimates of variance components and range by model (posterior mean ± standard deviation)

| Model | DIC | $\sigma_g^2$ | $\sigma_h^2$ | $\sigma_s^2$ | $\rho$ (km) | $\sigma_e^2$ |
| --- | --- | --- | --- | --- | --- | --- |
| G | 32716 | $0.60 \pm 0.02$ | - | - | - | $0.29 \pm 0.00$ |
| GH | 32650 | $0.14 \pm 0.01$ | $0.37 \pm 0.02$ | - | - | $0.29 \pm 0.00$ |
| GS | 32735 | $0.29 \pm 0.01$ | - | $0.32 \pm 0.06^a$ | $32.4 \pm 7.5$ | $0.29 \pm 0.00$ |
| GHS | 32596 | $0.11 \pm 0.01$ | $0.15 \pm 0.01$ | $0.32 \pm 0.05^b$ | $33.4 \pm 8.4$ | $0.29 \pm 0.00$ |

G - model with breeding value and residual, GH - model G plus herd effect, GS - model G plus spatial effect, and GHS - model GH plus spatial effect;
DIC - Deviance Information Criterion, $\sigma_g^2$ - genomic variance, $\sigma_h^2$ - herd variance, $\sigma_s^2$ - “realised” spatial variance, $\rho$ - spatial range (km), and $\sigma_e^2$ - Residual variance;
<sup>a</sup> - model parameter estimate was $0.50 \pm 0.12$ and <sup>b</sup> - model parameter estimate was $0.50 \pm 0.11$

The models showed important differences in estimated variance components as shown in Table 3. The estimate of genomic variance ranged substantially between the G model (0.60±0.02) and the GHS model (0.11±0.01). The other component of variance present in all models was the residual variance, which had a stable estimate (0.29±0.00) across all the models, suggesting that the difference between models was not due to different modelling of residual variation. The estimate of herd variance was larger in the GH model (0.37±0.02) compared to the GHS model (0.15±0.01), which is expected since the herd effect in GH model can capture some of the spatial effect. The estimate of “realised” spatial variance was similar between the GS model (0.32±0.06) and the GHS model (0.32±0.05). The estimate of spatial variance as a model parameter was larger than “realised” spatial variance between the herd location effects (0.50±0.12 for the GS model and 0.50±0.11 for the GHS model). The spatial range was on average larger in the GHS model (33.4±8.4 km) than in the GS model (32.4±7.5 km), but posterior distributions overlapped substantially. Attempting to estimate permanent environmental effects in the models for these data led to almost a complete “transfer” of genomic variance to permanent environmental variance, while estimates of herd and variance parameters were comparable (Supplementary 2). Taken together, these estimates demonstrate the challenge of separating genetic and environmental effects in LMIC smallholder settings.

### Correlations between breeding value, herd and spatial effects

Figure 5 shows the differences in EBV between GH and GHS models against the estimated spatial effects from the GHS model. The magnitude of the differences ranged from -0.4 to +0.4 and showed a moderate correlation (*R*^2^ = 0.57) with the estimated spatial effects. These results indicate that EBV from the GH model were overestimated for animals in good environments and underestimated for animals in worse environments compared to the GHS model.

**Figure 5.**
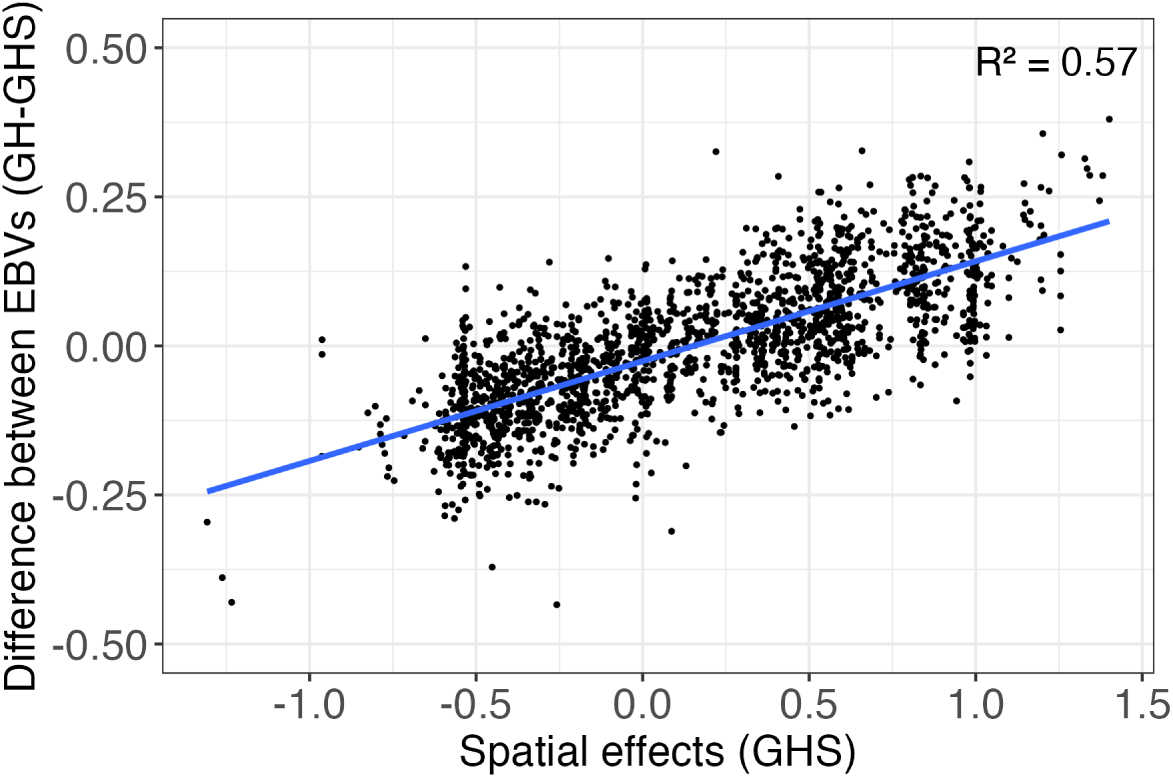
Difference in estimated breeding values (EBVs) between GH and GHS models against the estimated spatial effect from GHS model

The Table 4 shows Pearson’s rank correlations (upper diagonal) and Spearman’s rank correlations (lower diagonal) between estimated contribution of breeding values, herd effects, and spatial effects to phenotype values across models. There were substantial differences in correlations of breeding value contribution to phenotype values between models as already indicated in Figure 5. The Spearman’s correlation between breeding value contribution to phenotype values from G and GH models was 0.77, G and GS models was 0.74, and G and GHS models was 0.63. While such changes are expected when comparing with the G model, we also observed substantial differences in these correlations between the models GH, GS, and GHS. The Spearman’s correlation between breeding value contribution to phenotype values from GH and GS models was 0.65, GH and GHS models was 0.84, and GS and GHS models was 0.86.

**Table 4:** Pearson’s (above diagonal) and Spearman’s Rank (below diagonal) correlations between estimated contribution of breeding values (BV), herd effects, and spatial effects to phenotype values across models.

|  | BV_G | BV_GH | BV_GS | BV_GHS | Herd_GH | Herd_GHS | Spatial_GHS |
| --- | --- | --- | --- | --- | --- | --- | --- |
| BV_G | 1.00 | 0.78 | 0.76 | 0.66 | 0.92 | 0.66 | 0.64 |
| BV_GH | 0.77 | 1.00 | 0.68 | 0.87 | 0.52 | 0.33 | 0.43 |
| BV_GS | 0.74 | 0.65 | 1.00 | 0.87 | 0.65 | 0.87 | 0.06 |
| BV_GHS | 0.63 | 0.84 | 0.86 | 1.00 | 0.42 | 0.54 | 0.06 |
| Herd_GH | 0.92 | 0.53 | 0.62 | 0.40 | 1.00 | 0.72 | 0.71 |
| Herd_GHS | 0.64 | 0.32 | 0.85 | 0.53 | 0.69 | 1.00 | 0.06 |
| Spatial_GHS | 0.65 | 0.44 | 0.06 | 0.06 | 0.71 | 0.06 | 1.00 |

G - model with breeding value and residual, GH - model G plus herd effect, GS - model G plus spatial effect, and GHS - model GH plus spatial effect

Correlations between estimated contribution of breeding values, herd, and spatial effects to phenotype values showed that the models separated genetic and environmental effects differently (Table 4). For models G and GH, the Spearman’s correlation between estimated contribution of breeding value and herd effect to phenotype values (from model GH/GHS) was respectively 0.92/0.64 and 0.53/0.32. For models GS and GHS, it was respectively 0.62/0.85 and 0.40/0.53. These correlations show that all models produced EBV that were positively correlated with herd effect, but the magnitude of correlation ranged substantially. Even models that fitted the herd or spatial effect produced EBV that were correlated with herd effect, indicating true correlation or genetically better animals in better herds or insufficient data structure for better separation of these effects. For models G and GH, the Spearman’s correlation between estimated contribution of breeding value and spatial effect to phenotype values was respectively 0.65 and 0.44, whereas for models GS and GHS it was 0.06. These correlations show that EBV from G and GH models were capturing spatial effects, which was removed with the GS and GHS models. We also compared the top 50, 100, and 1000 ranked cows on EBVs from the GH and GHS models. For these three numbers of top cows we respectively observed an overlap of 35, 70, and 836] cows between the model (results not shown).

### Accuracy of phenotype prediction

Table 5 and Table 6 respectively present the cross-validation and forward-validation accuracy of phenotype prediction for the models. Spatial modelling increased the accuracy of phenotype prediction. In cross-validation by the percentage of exotic genome (predicting phenotypes of unobserved level based on other levels), we observed the highest average accuracy with GHS model (0.60), followed by GS model (0.56), GH model (0.40), and G model (0.33) (Table 5). This order was largely the same within each level of the percentage of exotic genome. Cross-validation accuracies were highest for the intermediate percentage of exotic genomes ((87.5, 60] and (60, 36]). In cross-validation by region (predicting phenotypes in an unobserved region based on other regions), all models showed low accuracy; GH model (0.16), G model (0.14), GS model (0.13), and GHS model (0.10) (Table 5). Cross-validation accuracies were lowest for the NE and NC regions.

**Table 5:** Cross-validation accuracy of phenotype prediction by model, percentage of exotic genome, and region.

| Percentage of exotic genome |  |  |  |  |  |
| --- | --- | --- | --- | --- | --- |
| Model | [100, 87.5]<br>(n=731) | (87.5, 60]<br>(n=770) | (60, 36]<br>(n=309) | (36, 0]<br>(n=84) | Average |
| G | 0.27 | 0.31 | 0.34 | 0.40 | 0.33 |
| GH | 0.36 | 0.45 | 0.45 | 0.35 | 0.40 |
| GS | 0.53 | 0.62 | 0.60 | 0.48 | 0.56 |
| GHS | 0.57 | 0.65 | 0.64 | 0.53 | 0.60 |

| Region |  |  |  |  |  |
| --- | --- | --- | --- | --- | --- |
| Model | NE<br>(n=437) | NC<br>(n=775) | SC<br>(n=414) | SW<br>(n=268) | Average |
| G | 0.06 | 0.07 | 0.27 | 0.15 | 0.14 |
| GH | 0.10 | 0.10 | 0.26 | 0.17 | 0.16 |
| GS | 0.02 | 0.01 | 0.24 | 0.15 | 0.13 |
| GHS | 0.11 | -0.02 | 0.18 | 0.14 | 0.10 |

G - model with breeding value and residual, GH - model G plus herd effect, GS - model G plus spatial effect, and GHS - model GH plus spatial effect
NE - North-East, NC - North-Central, SC - South-Central, and SW - South-West

**Table 6:** Forward validation accuracy of phenotype prediction by model, percentage of exotic genome, and region.

| Percentage of exotic genome |  |  |  |  |  |
| --- | --- | --- | --- | --- | --- |
| Model | [100, 87.5]<br>(n=60) | (87.5, 60]<br>(n=56) | (60, 36]<br>(n=24) | (36, 0]<br>(n=6) | Average |
| G | 0.29 | 0.28 | 0.54 | 0.83 | 0.49 |
| GH | 0.36 | 0.63 | 0.66 | 0.33 | 0.50 |
| GS | 0.37 | 0.59 | 0.73 | 0.74 | 0.61 |
| GHS | 0.44 | 0.67 | 0.75 | 0.66 | 0.63 |

| Region |  |  |  |  |  |
| --- | --- | --- | --- | --- | --- |
| Model | NE<br>(n=30) | NC<br>(n=60) | SC<br>(n=20) | SW<br>(n=36) | Average |
| G | 0.38 | 0.46 | 0.11 | 0.08 | 0.26 |
| GH | 0.65 | 0.57 | 0.32 | 0.35 | 0.47 |
| GS | 0.57 | 0.53 | 0.28 | 0.37 | 0.44 |
| GHS | 0.65 | 0.60 | 0.37 | 0.46 | 0.52 |

G - model with breeding value and residual, GH - model G plus herd effect, GS - model G plus spatial effect, and GHS - model GH plus spatial effect
NE - North-East, NC - North-Central, SC - South-Central, and SW - South-West

In forward validation by the percentage of exotic genome (predicting phenotypes of younger non-phenotyped animals based on older phenotyped animals), we observed the highest average accuracy with GHS model (0.63), followed closely by GS model (0.61), then GH model (0.50), and G model (0.49) (Table 6). As with cross-validation, predicting phenotypes of animals with highest percentage of exotic genome was the most challenging. In forward validation by region (predicting phenotypes of non-phenotyped animals based on older phenotyped animals), GHS model (0.52) performed best, followed closely by GH model (0.47), then GS model (0.44), and G model (0.26) (Table 6).

The spatial modelling increased accuracy as well in models with permanent environmental effect(GP models, see supplementary 3).

## Discussion

In this study, we evaluated the impact of leveraging precise herd location (GPS coordinates) on estimates of variance components, breeding values, and acurracy of phenotype prediction for milk yield in the LMIC smallholder setting. The exploratory data analysis showed a strong geographical context of the data in this study, including clustering of herds in four regions of Tanzania, the associated difference in mean milk yield between regions, covariation between the first few principal components of SNP genotypes between regions, and substantial variation within the regions. The estimated spatial effects expectedly showed large regional differences, and impacted estimates of variance components and breeding values. Furthermore, spatial modelling increased the accuracy of phenotype prediction, by establishing environmental connectedness between the herds, leading to a more accurate separation of genetic and environmental effects. These results highlight three broad points for discussion: (i) sources of spatial variation, (ii) spatial modelling improves estimates of environmental and genetic effects, and (iii) limitations of this study and future directions.

### Sources of spatial variation

The results show important differences in milk yield between regions. Lower milk yields in the North-East and South-Central regions of Tanzania, and higher milk yields in the North-Central and South-West regions of Tanzania. These differences are in line with the natural environmental conditions in Tanzania, specifically with altitude, precipitation, and temperature [31]. Furthermore, the North-East region is mountainous, including Mount Kilimanjaro and Mount Meru, characterised by low rainfalls and hence the negative effect on milk yield. Similarly, the South-Central region in our study corresponds to the Southern Highlands zone (e.g., Iringa region) with limiting conditions of high temperature (tropical) and humidity with moderate rainfall for producing sufficient quantity and quality of forage. On the other hand, the conditions in the North-Central region (e.g., Usa river and Themi River) and in South-West region (e.g., Kiwira River) characterised by high altitude and cooler with high amount of rainfall, impacting the flora and providing positive conditions for milk production.

### Spatial modelling improves estimates of environmental and genetic effects

Our results show that spatial modelling significantly affected model fit, estimated variance components and breeding values. Adding the spatial effect to the model always improved DIC, even when herd effect was already in the model. Also, estimate of genomic variance and herd variance changed with the addition of the spatial effect. Most importantly, the EBV from spatial models had practically no correlation with the estimated contribution of spatial effects to phenotype values, unlike the EBV from the standard genetic model with herd effect. The difference in EBV from the two models (GH and GHS) correlated positively with the estimated spatial effect, showing likely underestimation for animals in less favourable environments and overestimation for animals in more favourable environments. Correcting for such confounding is critical if we want to identify resilient animals that perform well in less favourable environments. These results indicate that spatial modelling better separated the shared environmental variation among nearby herds from other effects that contributed to phenotype variation. However, the EBV from both of these models correlated positively with the estimated contribution of herd effects to phenotype values. It is unclear from this study whether this positive correlation indicates another source of confounding or a true correlation whereby herds with better management have cows with higher breeding values; possibly due to admixture with exotic breeds.

Previous simulation study by Selle et al. [11] has shown that spatial modelling improves the separation of environmental and genetic effects. Like the genetic relationship/covariance matrix enables sharing information among relatives, spatial covariance matrix enables sharing information among nearby herds. In other words, spatial modelling encodes “expected” environmental connectedness among herds. We emphasise “expected” because this study used the Euclidean distance-based Matérn covariance function, without modelling environmental covariates at herd locations.

To assess potential improvements in EBV due to spatial modelling, we evaluated the accuracy of phenotype prediction in cross-validation and forward validation. Due to substantial admixture and regional differences, we validated the predictions by the percentage of exotic genome and region. In the previous analysis of the same data, Mrode et al. [4] assessed the accuracy by correlating EBV of cows with their yield deviations. However, in the present study we evaluated the accuracy of phenotype prediction, because of the challenging data structure. Namely, EBVs from a model without the spatial effect can contain such effects (as demonstrated in our study) and will hence better predict yield deviations, which also contain a part of the spatial effect not captured by the herd effect. In our study, spatial modelling always improved the accuracy of phenotype prediction in cross-validation and forward validation by the percentage of exotic genome. Cross-validation by region was challenging for all models, which is expected - all phenotype values in a region were masked and predicted based on a model trained on phenotype values from other regions. In forward validation by region, modelling spatial and herd effects jointly improved the accuracy of phenotype prediction beyond modelling just herd effect or just spatial effect, though the improvement of adding the spatial effect on top of the herd effect within each region was not large. While modelling the spatial effect seemed beneficial when looking at the overall model fit, estimated variance components and breeding values, we attribute this small improvement in forward validation to the small dataset and limited ways of being able to construct for a validation set.

We studied the use of spatial modelling for genetic evaluation, because many LMIC settings are characterised by very small herds (more than 93% of herds having less than 3 cows and more than 71% herds with a single cow in this study), which is an extremely challenging data structure for genetic evaluation. In this case, the standard GH model was not effective in separating genetic and environmental effects, because the herds were too small for their effect to be estimated accurately, leading to confounding between environmental and genetic effects. Several authors have studied the issue of small herd effects in genetic evaluation (summarised by [12]), including in the LMIC smallholder setting [3, 4, 5, 13, 18]. Chawala et al. [18] grouped herds into villages to fit them as a fixed effect. Powell et al. [5] found that modelling herd effects as random resulted in higher accuracy of EBV than modeling them as fixed when herd size is small (*<* 4 cows). Similarly, Costilla et al. [13] found that the accuracy of EBV with random herd effect improves as herd size increases, reaching its peak around five cows. Neither of these studies have analysed how their EBVs correlate with the estimated contribution of environmental effects to the observed phenotype values. We found three animal breeding studies that modelled environmental variation via spatial effects [11, 12, 15]. Sæbø and Frigessi [15] used the Besag’s model to account for large differences between veterinary districts, which is relevant for large differences in milk yield between regions in our study, but we also observed substantial variation in milk yield within regions. Selle et al. [11] used the same spatial modelling approach as in this study to analyse simulated data and sub- sampled real data from Slovenia (to mimic the LMIC setting), though predominant use of artificial insemination in the real data meant that there was more genetic connectedness than in most of LMIC smallholder settings. Cuyabano et al. [12] used exponential covariance function between beef herds in South Korea with the average herd size of 17 animals and more than 66% of farms having more than 5 animals, far more than in the LMIC smallholder setting. Compared to these studies, our results show promising improvement and hence expanded the past work on genetic evaluation in the LMIC smallholder setting [3, 4, 5, 13, 18].

### Limitations of this study and future directions

Our results indicate that spatial modelling can separate genetic and environmental effects more accurately compared to standard genetic models. However, to benefit from its full potential, breeders should ensure broad geographical coverage when recording data to maximise the accuracy of EBV, as shown with our cross-validation by region. The data used in this study are from the early stages of a quantitative genomic initiative for dairy cattle in Tanzania and hence had a limited coverage of regions in Tanzania as well as a limited number of herds per region. More data is being collected that will cover other regions in Tanzania, as well as other AADGG countries. To optimize limited phenotyping resources, we could allocate phenotyping based on variation between and within regions and the establishment of environmental connectedness with spatial modelling, possibly by extending the work of optimal selection of core individuals (see [32] and its references).

In this study we used distance-based covariance function for spatial effect, but increasingly more environmental data are available, such as temperature, precipitation, and soil quality, or other correlates [33]. These environmental covariates can be used to form “realised” environmental covariance matrix [34], like genomic markers (covariates) are used to form “realised” genomic covariance matrix [7]. When environmental covariates do not explain all the variance of spatial effects or are not available for all locations, the expected Euclidean distances and the realised distances could be combined like pedigree and genomic relationships are combined [8].

While spatial modelling improved model fit and other statistics in this study, we envision situations where analysts should be careful in fitting the spatial effect without validation. For example, migration and local adaptation due to selection can lead to a true correlation between environmental and genetic effects on phenotype values. In this setting, spatial effect will capture both environmental and genetic effects. Establishing genetic connectedness (see [35, 36] and their references) via designed mating plans is critical to enable separation of environmental and genetic effects. However, it is also important to understand processes that drive genetic variation across a geographical landscape (e.g., [37, 38, 39, 40, 41]).

For example, substantial genetic and environmental variation between and within regions, movements of cows between herds, and admixture with imported exotic germplasm are all typical in the LMIC smallholder setting, and having a deeper understanding of these processes could further improve modelling of genetic and environmental effects. In our study, we followed previous study of the same data and used their estimated percentage of exotic genome to model admixture of the studied animals [4]. Our principal component analysis showed that the first component correlated with these estimated percentages of exotic genome, as expected given the known admixture process. In addition, the second and third components differed significantly between some regions. For example, the North-East region of Tanzania has more challenging environmental conditions for milk production than other analysed regions, but also had a smaller percentage of exotic genome compared to other analysed regions. It is unclear if these differences are solely due to fewer imports of exotic animals/semen in the region or is natural selection in the region favouring animals with a smaller percentage of exotic genome. Further studies are required to address these topics, as well as topics related to modelling admixture involving multiple breeds of Taurine and Indicine origins, which challenge the accurate estimation of genetic effects due to different genomic history and associated linkage disequilibrium, possibly even non-additive genetic effects (e.g., [42, 43]).

The above example indicates a possible Genotype-by-Environment (GxE) interaction effect, which is likely to be important in the LMIC smallholder setting, especially when local and exotic breeds are used (e.g., [44, 45, 46]). Recently, Ducrocq et al. [46] indicated presence of GxE in their analysis of Holstein production in different regions of South Africa as different correlated traits or with a reaction norm involving herd climate characteristics. We did not consider GxE interactions in our study, due to a small dataset and already challenging data structure. However, given the wide range of environmental differences between Tanzanian regions and the significant cattle admixture in Tanzania, we anticipate that GxE analysis could be an important future focus. These future studies could test: i) how the effects of the SNP markers differ between Tanzanian regions, ii) how much GxE variance is there due to such effects, and iii) whether this is scale or crossover-type of GxE to understand possible re-ranking of animals when considering different environments.

Standard animal breeding software will require an extension to implement spatial modelling. We have used the R-INLA [25, 26] and inlabru [28], as these packages allow for a spatial effect as one of the model components. These packages use the same underlying linear algebra as standard animal breeding software, with extension to full Bayesian analysis [47]. Although R-INLA has been used in animal and plant breeding studies [11, 48, 49], it does not support all quantitative genetic models used in different breeding fields. Moreover, following a full Bayesian approach, an R-INLA user has to set prior distributions for all model parameters. Further developments allowing for the spatial effect in standard animal breeding software (e.g., ASReml, blupf90, MiX99) will be useful for the animal breeding community.

## Conclusions

The key insight of this study is that spatial modelling of herd effects more accurately separated genetic and environmental effects compared to standard models with independent herd effects, which in turn increases the accuracy of genetic evaluation. We have demonstrated these results with a milk yield dataset from smallholder farms of admixed cattle recorded as part of the Africa Asia Dairy Genetics Gains Programme in Tanzania. Nevertheless, we anticipate this structure of smallholder dairy systems is typical and similar across LMIC countries. Although the levels of proportion of exotic versus indigenous genomes in animals may differ across countries, the results of this study will guide future studies in other countries. Future studies are needed to explore the use of environmental covariates, the processes that drive environmental and genetic differences, including their interactions, and modification of standard animal breeding software to fit the spatial effect routinely. All these advances will enable better statistical modelling of the staggering variation in LMIC smallholder settings.

## Author’s contributions

IH and GG conceived the study. IH analysed the data and wrote the first draft of the manuscript. IP, GG, and RM provided statistical perspectives and reviewed the manuscript. GG supervised the study. Other authors have enabled the study through developing, supporting, and operating data recording system. All authors have read and approved the final manuscript.

## Availability of data and materials

The R scripts for this study are available in the Supplementary Material and at the GitHub repository https://github.com/HighlanderLab/ihouaga_adgg_spatial.

## Supporting information

Additional file 1 Posterior mean (left) and standard deviation (right) of the spatial effects from the GS model

Additional file 2 Estimates of variance components for models with permanent environmental effect

Additional file 3 Accuracy for models with permanent environmental effect

## Acknowledgements

The authors acknowledge funding from Royal Society Newton International Fellowship (NIF/R1/201150) and the Roslin Foundation grant (RF2303) to IH, BBSRC ISP grants (BBS/E/D/30002275, BBS/E/RL/230001A, and BBS/E/RL/230001C), Bill & Melinda Gates Foundation and UK aid from the UK Foreign, Commonwealth and Development Office (INV-040641) under the auspices of the Centre for Tropical Livestock Genetics and Health (CTLGH), established jointly by the University of Edinburgh, Scotland’s Rural College (SRUC), and the International Livestock Research Institute. The findings and conclusions contained within are those of the authors and do not necessarily reflect positions or policies of the Bill & Melinda Gates Foundation nor the UK Government.

## Authors note

For the purpose of open access, the authors have applied a CC BY public copyright license to any Author Accepted Manuscript version arising from this submission.

## Additional Files

Additional file 2 Estimates of variance components for models with permanent environmental effect

Additional file 3 Accuracy for models with permanent environmental effect

