## Additional file 1 Posterior mean (left) and standard deviation (right) of the spatial effects from the GS model for "Spatial modelling improves genomic evaluation in Tanzanian smallholder admixed dairy cattle"

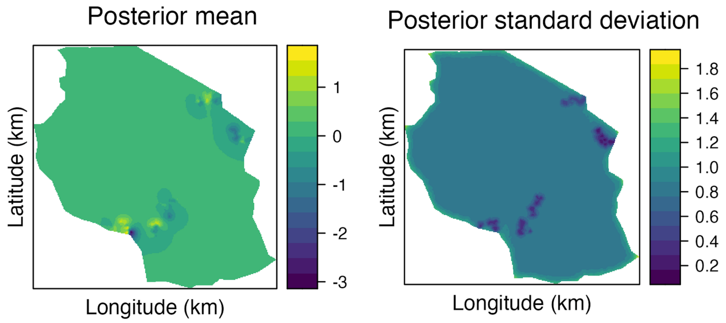


Figure S1: Posterior mean (left) and standard deviation (right) of the spatial effects (in units of phenotypic standard deviation) from the GS model
