## Additional file 2 Estimates of variance components for models with permanent environmental effect for "Spatial modelling improves genomic evaluation in Tanzanian smallholder admixed dairy cattle"

Table S2.1: Estimates of variance components and range by model, including permanent environment effect (posterior mean ± standard deviation)

| Model | DIC | $\sigma_g^2$ | $\sigma_h^2$ | $\sigma_{pe}^2$ | $\sigma_s^2$ | $\rho$ (km) | $\sigma_e^2$ |
|---|---|---|---|---|---|---|---|
| GP | 32692 | 0.03 ± 0.02 | - | 0.46 ± 0.03 | - | - | 0.29 ± 0.00 |
| GPH | 32633 | 0.02 ± 0.00 | 0.36 ± 0.02 | 0.12 ± 0.01 | - | - | 0.29 ± 0.00 |
| GPS | 32620 | 0.01 ± 0.01 | - | 0.23 ± 0.01 | 0.34 ± 0.07[a] | 33.5 ± 6.6 | 0.29 ± 0.00 |
| GPHS | 32557 | 0.01 ± 0.01 | 0.13 ± 0.01 | 0.11 ± 0.01 | 0.33 ± 0.06[b] | 32.3 ± 7.8 | 0.29 ± 0.00 |

GP - model with breeding value, permanent environment effect and residual, GPH - model GP plus herd effect, GPS - model GP plus spatial effect, and GPHS - model GPH plus spatial effect;
DIC - Deviance Information Criterion, $\sigma_g^2$ - genomic variance, $\sigma_h^2$ - herd variance, $\sigma_{pe}^2$ - permanent environment variance, $\sigma_s^2$ - "realised" spatial variance, $\rho$ - spatial range (km), $\sigma_e^2$ - Residual variance;
[a] - model parameter estimate was 0.57±0.14 and [b] - model parameter estimate was 0.52±0.12
