## Additional file 3 Accuracy for models with permanent environmental effect for "Spatial modelling improves genomic evaluation in Tanzanian smallholder admixed dairy cattle"

Table S3.1: Cross-validation accuracy of phenotype prediction by model, percentage of exotic genome, and region

| | Percentage of exotic genome | | | | |
|---|---|---|---|---|---|
| Model | [100, 87.5] (n=731) | (87.5, 60] (n=770) | (60, 36] (n=309) | (36, 0] (n=84) | Average |
| GP | 0.29 | 0.35 | 0.39 | 0.40 | 0.36 |
| GPH | 0.36 | 0.45 | 0.46 | 0.38 | 0.41 |
| GPS | 0.56 | 0.65 | 0.64 | 0.57 | 0.60 |
| GPHS | 0.58 | 0.67 | 0.66 | 0.57 | 0.62 |
| | Region | | | | |
| Model | NE (n=437) | NC (n=775) | SC (n=414) | SW (n=268) | Average |
| GP | 0.08 | -0.01 | 0.27 | 0.15 | 0.12 |
| GPH | 0.03 | 0.06 | 0.27 | 0.19 | 0.14 |
| GPS | 0.10 | -0.08 | 0.27 | 0.17 | 0.12 |
| GPHS | 0.04 | -0.03 | 0.16 | 0.13 | 0.08 |

GP - model with breeding value, permanent environmental effect and residual, GPH - model GP plus herd effect, GPS - model GP plus spatial effect, and GPHS - model GPH plus spatial effect

NE - North-East, NC - North-Central, SC - South-Central, and SW - South-West

Table S3.2: Forward validation accuracy of phenotype prediction by model, percentage of exotic genome, and region

| | Percentage of exotic genome | | | | |
|---|---|---|---|---|---|
| Model | [100, 87.5] (n=60) | (87.5, 60] (n=56) | (60, 36] (n=24) | (36, 0] (n=6) | Average |
| GP | 0.36 | 0.36 | 0.60 | 0.70 | 0.50 |
| GPH | 0.43 | 0.65 | 0.68 | 0.39 | 0.54 |
| GPS | 0.40 | 0.64 | 0.74 | 0.72 | 0.62 |
| GPHS | 0.48 | 0.70 | 0.75 | 0.76 | 0.67 |

| | Region | | | | |
|---|---|---|---|---|---|
| Model | NE (n=30) | NC (n=60) | SC (n=20) | SW (n=36) | Average |
| GP | 0.45 | 0.51 | 0.17 | 0.13 | 0.31 |
| GPH | 0.65 | 0.63 | 0.36 | 0.35 | 0.50 |
| GPS | 0.61 | 0.56 | 0.41 | 0.38 | 0.49 |
| GPHS | 0.66 | 0.63 | 0.44 | 0.47 | 0.55 |

GP - model with breeding value, permanent environmental effect and residual, GPH - model GP plus herd effect, GPS - model GP plus spatial effect, and GPHS - model GPH plus spatial effect

NE - North-East, NC - North-Central, SC - South-Central, and SW - South-West
